# Hybrid epigenomes reveal extensive local genetic changes to chromatin accessibility contribute to divergence in embryonic gene expression between species

**DOI:** 10.1101/2023.01.04.522781

**Authors:** Hannah R. Devens, Phillip L. Davidson, Maria Byrne, Gregory A. Wray

## Abstract

Chromatin accessibility plays an important role in shaping gene expression patterns across development and evolution; however, little is known about the genetic and molecular mechanisms that influence chromatin configuration itself. Because *cis* and *trans* influences can both theoretically influence the accessibility of the epigenome, we sought to better characterize the role that both of these mechanisms play in altering chromatin accessibility in two closely related sea urchin species. Using hybrids of the two species, and adapting a statistical framework previously developed for the analysis of *cis* and *trans* influences on the transcriptome, we examined how these mechanisms shape the regulatory landscape at three important developmental stages, and compared our results to similar patterns in the transcriptome. We found extensive *cis*- and *trans*-based influences on evolutionary changes in chromatin, with *cis* effects slightly more numerous and larger in effect. Genetic mechanisms influencing gene expression and chromatin configuration are correlated, but differ in several important ways. Maternal influences also appear to have more of an effect on chromatin accessibility than on gene expression, persisting well past the maternal-to-zygotic transition. Furthermore, chromatin accessibility near GRN genes appears to be regulated differently than the rest of the epigenome, and indicates that *trans* factors may play an outsized role in the configuration of chromatin near these genes. Together, our results represent the first attempt to quantify *cis* and *trans* influences on evolutionary divergence in chromatin configuration in an outbred natural study system, and suggest that the regulation of chromatin is more genetically complex than was previously appreciated.

## INTRODUCTION

Chromatin configuration plays a critical role in transcriptional regulation in eukaryotes by enabling the ability of regulatory elements to influence transcription (Kornberg and Lorch 1992; Giacoman-Lozano, et al. 2022). Over the past decade, genome-wide assays (Zaret 2005; Song and Crawford 2010; Buenrostro, Wu, Chang, et al. 2015; Cusanovich, et al. 2018) have revealed that chromatin accessibility is both highly dynamic and highly context-dependent. Extensive changes in the complement of open chromatin regions (OCRs) take place during development (Cusanovich, et al. 2018; Reddington, et al. 2020), leading to fully differentiated cells that typically differ in the majority of OCRs (Yue, et al. 2014; Buenrostro, Wu, Litzenburger, et al. 2015). Further changes in the accessibility of individual OCRs take place across circadian cycles and in response to a wide range of physiological conditions and external stimuli. The sheer scale of this remodeling is enormous: of the >800,000 OCRs known in humans, for instance, the vast majority are only accessible under a few conditions or in a small number of cell types or developmental stages (Thurman, et al. 2012; Jiang, et al. 2022).

These findings underscore that chromatin remodeling is an important mechanism contributing to transcriptional regulation, but the evolutionary significance of this remodeling remains poorly understood. The mechanistic basis for differences in chromatin accessibility can be thought of as occurring in two different ways: through changes in “*cis*” (changes to the nucleotide sequence of *cis*-regulatory elements themselves), or through changes in “*trans*” (alterations to the structure, localization, or expression of the transcription factors that interact with *cis*-regulatory elements) (see Figure 1). However, the relative contribution of *cis* versus *trans* changes to chromatin accessibility differences remains unclear. Understanding the role that each of these mechanisms play in changing chromatin accessibility is crucial to our understanding of how chromatin accessibility evolves, because they represent two fundamentally different explanations for alterations to chromatin status: while *trans* changes may occur simply as an indirect byproduct of an alteration to an upstream factor in response to a changed nuclear environment, *cis* changes reflect a direct influence of evolution on the sequence of the regulatory element itself. Thus, it is important to establish whether evolutionary differences in OCRs are heritable and consequential or merely a reflection of other genetic changes that instead influence transcription.

**Figure 1.**
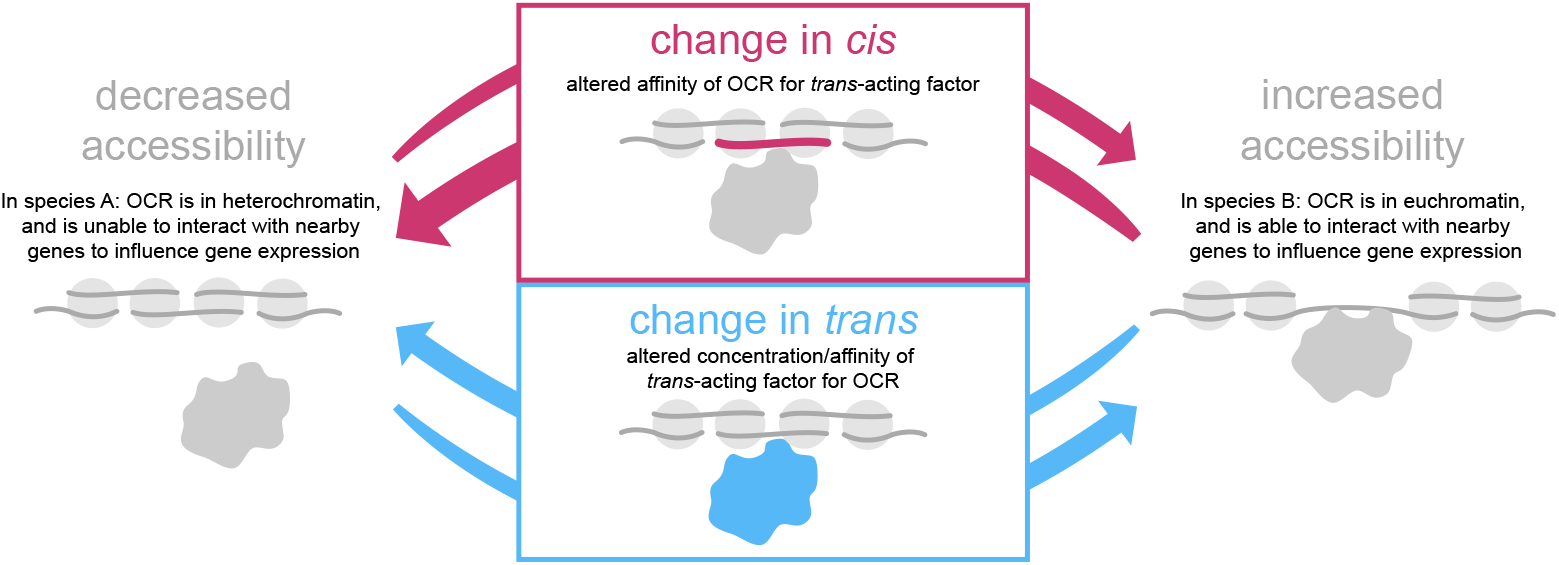
Conceptual overview of molecular mechanisms that could produce differences in chromatin accessibility between species. While several distinct molecular processes can modulate chromatin configuration, these can be grouped into two broad categories: those that are genetically based near the open chromatin region (OCR) of interest (*cis*) and those based elsewhere in the genome (*trans*). *Cis*-based changes (magenta here and in subsequent figures) are caused by a local mutation that alters binding of an already-present protein. Depending on whether the mutation raises or lowers binding affinity, and on the protein’s biochemical function, the consequence could be either an increase or decrease in accessibility at a specific genomic location (bidirectional arrows). *Trans*-based changes (light blue here and in subsequent figures) can be caused by a mutation that either alters the amino acid sequence or post-translational processing of a protein and thereby modifies its function, or by a change in the presence or concentration of a protein in the nucleus. (Note: elsewhere in this text we also refer to the nature and concentration of proteins available to interact with chromatin as the “nuclear environment”). Again, depending on the specific nature of these changes, chromatin accessibility at a specific location in the genome could either increase or decrease (bidirectional arrows).

Several studies have compared the open chromatin landscape among related species, in which it is possible to identify orthologous noncoding regions of the genome with high confidence (Shibata, et al. 2012; Pizzollo, et al. 2018; Edsall, et al. 2019; Lewis and Reed 2019; Swain-Lenz, et al. 2019; Davidson, et al. 2022; Yao, et al. 2022). These studies demonstrate that most OCRs are conserved in position and degree of accessibility among closely related species (<10 million years diverged), while a smaller proportion are conserved over longer time frames (Yue, et al. 2014; Gao, et al. 2018; Davidson, et al. 2022). These studies also reveal that hundreds or even thousands of OCRs are differentially accessible in the same cell type or tissue under the same conditions, even among closely related species. Some studies report a statistical association between whether an OCR is differentially accessible (DA) and whether the nearest gene is differentially expressed (DE) between species (Pizzollo, et al. 2018; Davidson, et al. 2022). This finding suggests that evolutionary changes in chromatin configuration may contribute to divergence in gene expression. That said, the association is generally weak and does not provide information about individual OCRs and genes. More importantly, it remains unclear to what extent evolutionary differences in chromatin status are heritable and how often heritable differences influence gene expression.

In this study, we used interspecies hybrids to measure genetic contributions to divergence in chromatin configuration and the relationship of chromatin configuration to gene expression during embryonic development. We took advantage of a “natural experiment” in the evolution of gene expression driven by a recent life history switch in sea urchins from planktotrophy (feeding larvae) to lecithotrophy (nonfeeding larvae) (Raff and Byrne 2006; Wray 2022). This life history switch produced an unusually high concentration of recent evolutionary changes in gene expression (Israel, et al. 2016) and chromatin configuration (Davidson, et al. 2022) on the branch leading to lecithotrophy. Positive selection is enriched on this branch, providing the ability to contrast neutral and adaptive changes in gene regulation (Israel, et al. 2016; Davidson, et al. 2022; Wray 2022). The evolution of lecithotrophy also involved extensive changes in maternal provisioning of metabolites and informational molecules (Hoegh-Guldberg and Emlet 1997; Byrne, et al. 1999; Israel, et al. 2016; Davidson, et al. 2019). These recent, extensive changes in the molecular composition of eggs also allows us to test whether divergence in chromatin configuration is associated with changes in gene expression or is simply an indirect consequence of altered physiology with little relevance for the evolution of gene expression. Furthermore, the well-defined developmental gene regulatory network (dGRN) of sea urchins (Davidson, et al. 2002; Oliveri, et al. 2008; Su, et al. 2009; Peter and Davidson 2011; Rafiq, et al. 2012) affords the opportunity to examine the architecture of *trans* effects in detail and to examine how natural selection operates on the transcriptional regulation of critical developmental genes.

We compared the open chromatin landscape during embryonic development in hybrids with those of same-species parental crosses, adapting a well-established statistical framework for analysis of hybrid transcriptomes (Wittkopp, et al. 2008; McManus, et al. 2010; Pirinen, et al. 2015) to analysis of hybrid epigenomes. Taken together, our results emphasize the difference in the regulation of the epigenome relative to the regulation of gene expression, support the idea that genetically based changes in chromatin contribute to evolutionary divergence of gene expression, and highlight several distinct evolutionary properties of OCRs near dGRN genes.

## METHODS

### Experimental design, animal husbandry, and sample processing

We generated hybrid embryos from *H. erythrogramma* females and *H. tuberculata* males as well as both same-species crosses (Figure 2A). (The reciprocal cross arrests as gastrulae (Raff, et al. 1999), so hybrids were generated in one direction only.) We made three biological replicates of each cross using independent parents for each set of replicate crosses. From these crosses we collected embryos at three developmental stages (blastula, gastrula, larva), matching those of our previous analysis of hybrid transcriptomes (Wang, et al. 2020) and a subset of stages in our previous comparative ATAC-seq study (Davidson, et al. 2022). We prepared ATAC-seq libraries from each of the 29 samples and generated 75b paired-end reads. Reads were aligned to reference genomes, yielding 3,850,031-40,309,001 mapped reads per sample (see Table S1, Table S3). We used macs2 (Zhang, et al. 2008)to identify transposase-accessible sites with an FDR of 5%, most of which are shared among species and present in hybrids (Figure 2D).

**Figure 2.**
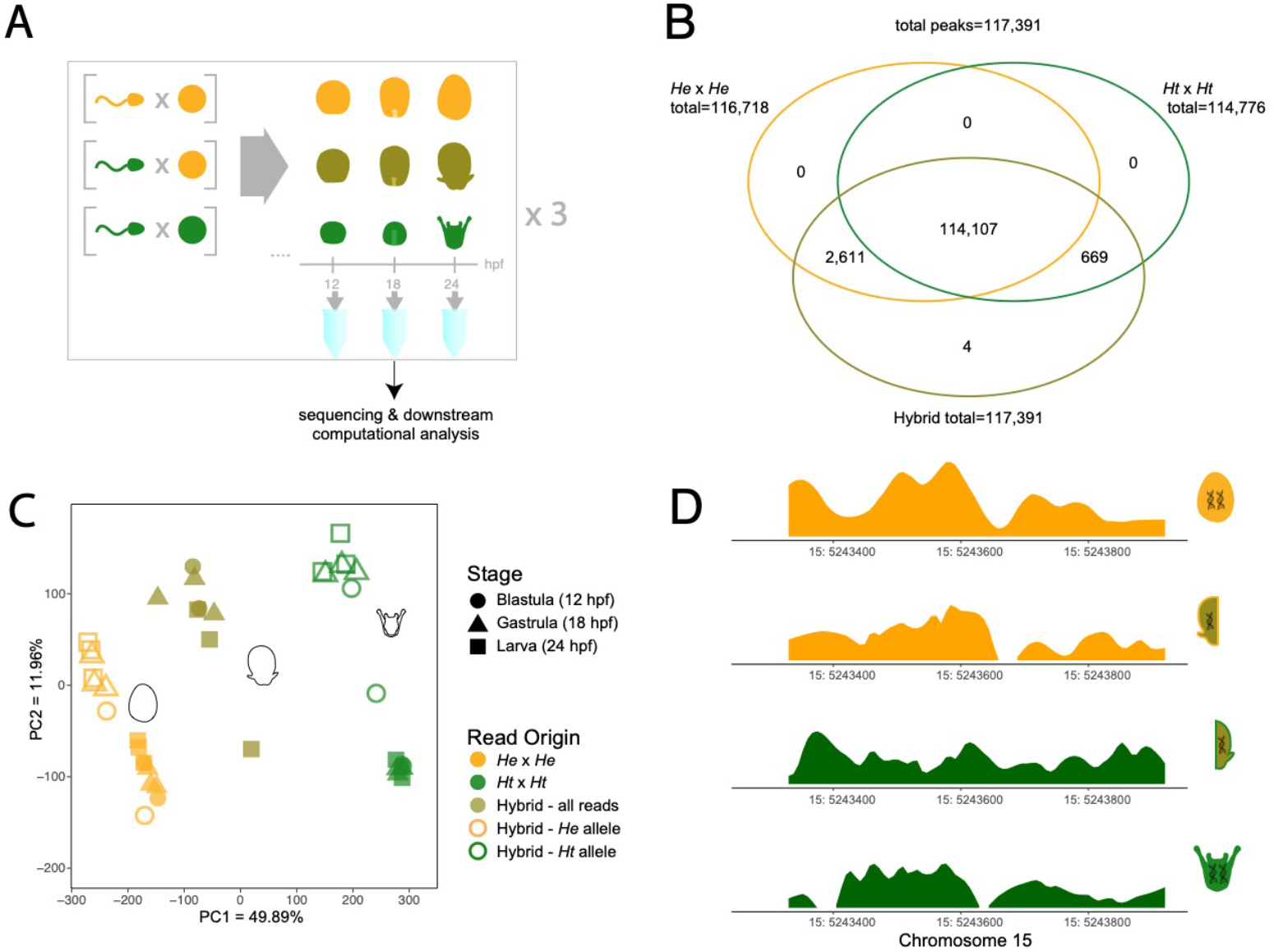
Chromatin configuration in parents and hybrids. **(A)** Experimental design and workflow. Samples from three biological replicates of three genetic crosses (*He* x *He*, the maternal same-species cross; *Ht* x *Ht*, the paternal same-species cross; and *He* (female) x *Ht* (male), the hybrid cross) were collected at three timepoints (12, 18, 24 hpf: hours post fertilization). **(B)** Venn diagram (not area-proportional) of peaks that are unique *vs*. shared among the same-species crosses and the hybrid cross. The reported peak count is the number of peaks following low-count removal. **(C)** Principal component analysis (PCA) of ATAC-seq results generated from counts table of reads in open chromatin regions. Throughout this study, orange indicates *He* origin; green *Ht* origin; and olive hybrid origin. **(D)** Real example of browser track for a peak with “conserved” regulation, indicating no statistically significant difference in accessibility for any of the crosses and, in the case of the hybrid, no statistically significant difference in the accessibility of either allele compared to the accessibility of the respective same-species cross. Note that for the same-species crosses, total accessibility is shown, while for the hybrid, the browser track has been broken out into the accessibility of the two different alleles.

Fertile *H. erythrogramma* and *H. tuberculata* adults were acquired from wild populations near Sydney, Australia and held in ^~^22 °C aquaria at the Sydney Institute of Marine Sciences. Aquaria were supplied with flow through ambient seawater. Cultures were produced from eggs and sperm obtained from these adults by intracoelomic injection of 0.5 KCl. A breeding design with three biological replicates was employed for each of the following crosse*s: H. erythrogramma* ♀ x *H. erythrogramma* ♂, *H. tuberculata* ♀ x *H. tuberculata* ♂, *H. erythrogramma* ♀ x *H. tuberculata* ♂ (Figure 2A). Cultures were fertilized and reared as previously described (Israel, et al. 2016; Wang, et al. 2020). Embryos were collected for analysis at three stages (blastula (12 hpf), gastrula (18 hpf), and larva (24 hpf)) and subjected to a modified version of the Omni ATAC-seq protocol(Corces, et al. 2017; Davidson, et al. 2022).

Hybrid sea urchins were generated in much the same manner as described previously (Wang, et al. 2020). Briefly, *H. erythrogramma* eggs were washed in acidified sea water (pH 5) for 60 seconds to remove the jelly coat and then washed twice in FASW. They were fertilized with excess *H. tuberculata* sperm and then washed twice in FASW. Cultures were grown at 22-24 C with daily water changes. Because fertilization rates using this method were low (around 5%), embryos were hand-picked at the time of collection (blastula = 50, gastrula = 50, larva = 5 embryos), yielding ^~^70,000 nuclei per sample.

The MinElute reaction cleanup kit (Qiagen) was used for sample purification, followed by library preparation using the Qiaquick PCR purification kit (Qiagen) and size selection using AMPure XP beads (Beckman Coulter). Reads were sequenced on the Illumina HiSeq 4000 platform at the Duke Center for Genomic and Computational Biology. 150 base-pair paired-end sequencing was used for the hybrid samples, while same-species samples were sequenced using a mix of 50 base-pair paired-end and single-end sequencing (see *Methods* of Davidson and Israel et al. 2022). In the interest of consistency across samples, the R2s of paired-end samples were discarded and all reads were analyzed as single-end; furthermore, all reads were trimmed to 50 bp in the trimming step. Raw reads were trimmed using TrimGalore (Krueger 2016) and the following parameter: trim_galore -q 20 --length 50 --fastqc. Next, reads were aligned to both *Heliocidaris* genomes using BBSplit, a read-binning aligner based on BBMap (sourceforge.net/projects/bbmap). Briefly, BBSplit uses BBMap to align a read to two genomes simultaneously, scores the alignments for mismatches, and retains the alignment with the higher score. This allows for each read in a sample to be assigned a parental genome of origin. As proof of principle (as well as for the sake of consistency in approach), all samples from same-species crosses were also aligned using BBSplit, with over 94% of each same-species cross’s reads mapping back to the “correct” parent-of-origin (Table S1). We also created “in silico” hybrid samples as another method of testing the ability of the BBSplit tool to correctly identify the genomic origin of a read. Briefly, we aligned samples from same-species crosses to their correct genome using BBMap, subsampled a given number of aligned reads from each same-species cross, concatenated these files together, and then re-aligned the resulting “hybrid” sample using BBsplit. If BBsplit were perfectly able to differentiate reads from the two parent genomes, 50% of these in silico hybrids should have mapped to *H. erythrogramma* and 50% should have mapped to *H. tuberculata*. In actuality, an average of 50.7% mapped to *H. erythrogramma* and 47.0% mapped to *H. tuberculata* (the remaining ^~^2% of reads could not be assigned confidently to one genome or the other). Breakdowns of parent-of-origin for the in-silico hybrids, as well as for each real hybrid sample, are available in Tables S2 and S3.

Reads were quality-filtered using SamTools (Danecek, et al. 2021) and reciprocal liftovers between the two reference genomes were performed using the UCSC LiftOver tool (Hinrichs, et al. 2006). Briefly, reciprocal liftovers allow for all sequences to be transposed into the coordinates of just one species’ genome (in this case, *H. erythrogramma*) while minimizing the reference bias that occurs when lifting coordinates between genomes. For *H. tuberculata* reads, alignments were lifted from *H. tuberculata* coordinates to *H. erythrogramma* coordinates with a -minMatch value of 0.5; for *H. erythrogramma* reads, alignments were lifted to *H. tuberculata* coordinates and then back to H. erythrogramma coordinates, all with a -minMatch value of 0.5. For all samples, only reads reciprocally lifting over to the *Heliocidaris erythrogramma* genome were retained, and *Heliocidaris erythrogramma* genomic coordinates were used for all further analysis. Peaks were called for each stage across all samples from the same species (*He, Ht* or hybrid) using macs2. Duplicate tags were also removed in the same step. The resulting narrowPeak files were combined and peaks were merged to create a master bed file, which allowed the generation of a counts table across all samples using the bedtools multicov function (Quinlan and Hall 2010). This counts table was the basis for further computational analyses conducted in R (version 4.0.2).

### Calculation of FRIP scores

A standard quality control metric for ATAC-seq studies is the fraction of reads in peaks (FRIP). We calculated FRIP scores on individual samples based on established ENCODE methods (https://www.encodeproject.org/data-standards/terms/#library) using reciprocally-lifted reads. However, a complication arises in dealing with reads from hybrid samples that were aligned to two different genomes. Since there is not (to our knowledge) a published approach for calculating FRIP scores with such reads, we opted to simply bin each hybrid sample into two “subsamples” based on the parental genome they best aligned to as described above; this approach resulted in two FRIP scores per hybrid sample. This method resulted in an average FRIP score of 23% for the hybrid crosses and 34% for the same-species crosses. While the FRIP scores for the hybrid crosses were lower than for the same species crosses, they were still “acceptable” according to ENCODE guidelines; moreover, the fact that these FRIP scores were obtained on essentially “half” samples suggests that the quality of reads in biologically relevant peaks was comparable for both the hybrids and the same-species crosses despite much lower read depth in the hybrid crosses. We also observed a vanishingly small number of “underdominant” peaks (see Results), which we would expect to be more common if the quality of our hybrid dataset remained poorer than that of our same-species dataset after our filtering steps (creation of a union peak set and removal of low-count reads) were performed.

### R analysis

Two extra samples from the *H. tuberculata* crosses (blastula stage and gastrula stage), which had been used to build a more complete peak set, were removed from the analysis for the purposes of balancing statistical power across stages. Furthermore, after initial PCA analysis, it was determined that one of the *H. erythrogramma* blastula samples was an outlier (potentially due to its small library size), and thus this sample was also removed (along with a corresponding *H. tuberculata* sample to ensure a balance in statistical power of downstream tests). This left a total of 26 samples (though for some analyses the hybrid samples were split in “half” when reads were aligned to their inferred parent of origin, leading to a total of 34 “samples”--see Table S5). After low-count removal using R’s *cpm* function, a total of 117,391 peaks remained. Read counts for each peak were *vst*-transformed, and principal component analysis (PCA) was completed on these *trans*formed reads using the R *prcomp* function. Inheritance and regulatory modes were defined and calculated as described in (Coolon, et al. 2014; Wang, et al. 2020)(see Table S6 for classification parameters).

Differential accessibility analysis was performed using the DESeq2 package (Love, et al. 2014) in R. Mean expression change distributions, based on expression values from (Israel, et al. 2016) were compared using one-way ANOVAs and Scheffe’s Test. These tests work for datasets which may have unequal variances, so they are appropriate for comparing distributions with large differences in the number of observations per dataset (as was the case for many of our comparisons). The GRN gene set used was the same as in a previous analysis (Davidson, et al. 2022), and was originally obtained from BioTapestry.org. Gene functional categories were obtained from Echinobase (www.echinobase.org).

## RESULTS

### Reads from hybrid embryos reflect expected biological signals, including reproducing parent-of-origin patterns

We generated ATAC-seq libraries from maternal *Heliocidaris erythrogramma* (*He*) x paternal *H. tuberculata* (*Ht*) hybrid embryos from three independent crosses at three stages of development (Figure 2A) and assigned reads to parental genomes (see Methods). We refer to these crosses, respectively, as either “hybrids” or “same-species” throughout. The stages sampled match those in our earlier analysis of transcriptomes in the same hybrid cross (Wang, et al. 2020) and are a subset of stages examined in comparative transcriptome and epigenome time courses for the two *Heliocidaris* species and an outgroup, *Lytechinus variegatus* (Israel, et al. 2016; Davidson, et al. 2022). We present data here from only one direction of interspecies hybrids because the reverse cross (maternal *Ht* x paternal *He*) arrests during gastrulation (Raff, et al. 1999). Before quality filtering, an average of 92.2% of reads from hybrids could be confidently assigned to a parental genome using this workflow (see Methods, Table S3). This is a marked improvement over the approach we previously used in the transcriptome, which was able to map only 81.9% of reads to a parental genome. On average, 56.9% of our hybrid reads mapped to the *H. erythrogramma* genome and the remaining 43.1% mapped to the *H. tuberculata* genome (Table S3). After quality filtering, we calculated FRIP scores as described in Methods. Of reciprocally lifted hybrid reads aligned to the *H. erythrogramma* genome, an average of 29.8% fell in peaks called on hybrids, while an average of 16.3% of hybrid reads aligned to the *H. tuberculata* genome fell in peaks called on hybrids.

We identified a total of 124,282 OCRs in hybrids across the three developmental stages examined. Of these, 117,391 OCRs (94.5%) occur within regions of the genome that are 1:1 orthologous in the two parental genomes. A further 6,891 OCRs (5.5%) occur within regions that are entirely absent in the genome of one or the other parental species. In subsequent analyses we refer to these as orthologous and paralogous sets of OCRs, respectively. The overwhelming majority of orthologous OCRs in hybrids (117,387 or >99.999%) fall within OCRs called on same-species crosses, with just 4 OCRs present only in hybrids (Figure 2B). Among the set of orthologous OCRs, 37,010 (31.5%) are differentially accessible (DA) between species at one or more developmental stages. The proportion of DA OCRs at each developmental stage parallels that in our previous study despite the sets of OCRs being independently called (i.e., they overlap but are not identical). The high degree of concordance in peak calling and parallel fractions of OCRs identified as DA indicates that the ATACseq data from hybrids reported here is comparable in quality to our published data from the same-species crosses (Davidson, et al. 2022).

As a further check on data quality, we examined peak height in our dataset. Reads in hybrids that map to putative promoter regions (the first peak within 500 bp upstream of the translation start site, TLS) formed more open peaks than those that map to putative distal enhancers (those >500 bp from the nearest translation start site) (Figures S1, S2). This result is consistent with many other studies that find core promoter regions to be generally more open than distal enhancers (Klemm, et al. 2019). Peaks in hybrids corresponding to putative promoter regions and enhancer regions are similar in size to those in same-species crosses (Davidson, et al. 2022), suggesting that biological effects outweigh technical influences on read mapping in hybrids.

To more formally assess the primary drivers of differences in ATAC-seq reads among samples, we carried out Principal Component Analysis of the hybrid and same-species data (Figure 2C). Principal component (PC) 1 explained 49.89% of the variation, separating samples by species, while PC2 explained a further 11.96% of the variation, separating the hybrid reads from the same-species crosses. Along PC1, the combined hybrid reads for each sample (i.e., the sum of reads from both chromosomes) fell approximately halfway between samples from same-species crosses (Figure 2C, solid symbols). These results indicate that both parents contribute to variation among samples. Given that chromosomes from both parents are exposed to a common molecular environment in hybrids, this result further suggests that parental genotype has a substantial influence on chromatin configuration. When reads from hybrids were separated by inferred parent-of-origin, however, each sample clustered much closer to samples from the matching same-species crosses (Figure 2C, open symbols). This result indicates that assignment of individual reads to parent-of-origin is generally correct. Note that a perfect overlap with the matching same-species samples is not necessarily expected even if every read is accurately assigned, as this would only occur in the absence of any *trans* genetic effects (i.e., if the molecular environment of hybrid nuclei had no influence on chromatin different from that in the same-species cross).

None of the first four PCs separate reads by developmental stage (Figures 2C, S3), suggesting that the epigenome as a whole does not change extensively across the three stages sampled in the *Heliocidaris* species or their hybrids. Stage-to-stage differential accessibility analysis confirmed this finding in each cross, as fewer than 0.06% of peaks were differentially accessible between stages in all crosses studied (p<=0.1) (Table S7). This result is in contrast to the transcriptome, where PC1 separates samples by developmental time across the same three stages examined here (Israel, et al. 2016; Wang, et al. 2020)(Figure 2C).

### Mapping hybrid reads to parent-of-origin reveals distinct genetic effects

The PCA results imply that differences in OCRs between the two *Heliocidaris* species have a substantial genetic component. To explore the genetic basis for divergence in chromatin configuration among species in more detail, we adapted a statistical framework originally developed for hybrid transcriptomes (Coolon, et al. 2014) and applied it to the set of orthologous OCRs. This approach uses a series of statistical tests to classify the genetic basis for a quantitative trait, in this case normalized ATACseq read counts, in terms of inheritance mode (dominance effects) and regulatory mode (*cis* and *trans* effects) (Table S6). The majority of orthologous OCRs (68.5%) are not differentially accessible between species at any of the three stages of development we examined (as in (Davidson, et al. 2022)) and are thus classified as conserved (see Figure 2D for an example of a “conserved” OCR). The fraction of OCRs with conserved chromatin status is highest at blastula, the earliest stage examined (Figure 3A, B). This is notably different from the transcriptome, which becomes progressively more similar between the two *Heliocidaris* species during development (Israel, et al. 2016; Wang, et al. 2020).

**Figure 3.**
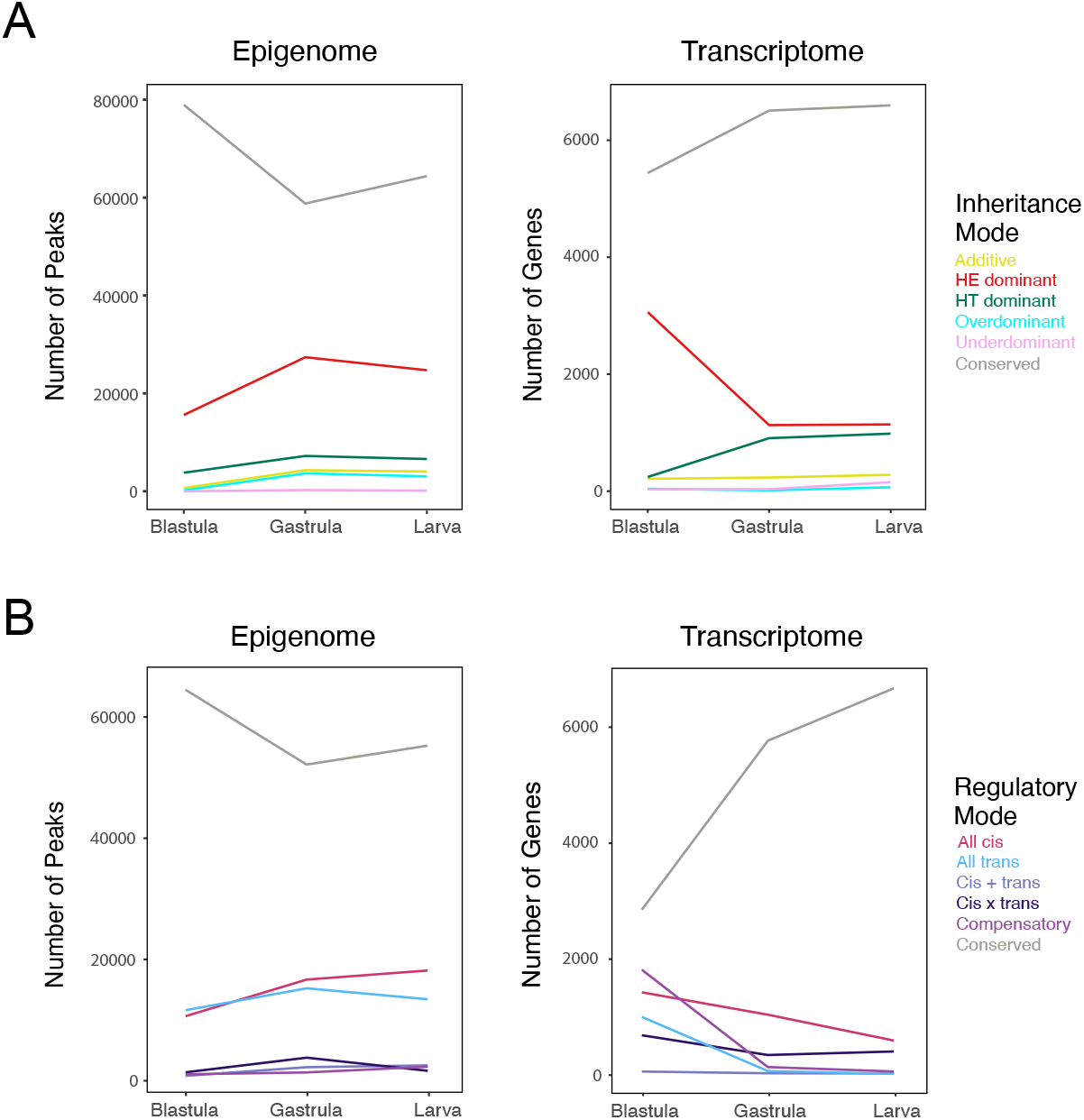
Contrasts between genetic basis for evolutionary changes in chromatin configuration and in transcript abundance. **(A)** Line plots of inheritance mode classification for all open chromatin regions (left) and for all genes (right). **(B)** Line plots of regulatory mode classification for all open chromatin regions (left) and for all genes (right). Transcript abundance data from Wang et. al (2020).

Of the 37,010 orthologous OCRs that differ in accessibility between species, from 84.4-87.6% could be classified according to inheritance mode, depending on developmental stage (Figure 3A, left). At all stages, the majority of these are inferred to be simple dominance effects, with just a small number of additive, underdominant, and overdominant effects. Note that the four OCRs present only in hybrids (mentioned previously) are extreme examples of overdominance. We also sought to classify the set of DA OCRs by regulatory mode, and found that 76.6-79.5% of the orthologous OCRs that differ in accessibility between species could be classified by regulatory mode (Figure 3B, left). The majority of these are inferred to be all-*cis* or all-*trans* effects (see Figure 4 for a visual explanation and examples of browser tracks for *cis* and *trans* effects). A very small fraction of differential OCRs are inferred to reflect various forms of *cis-trans* interactions (*cis* x *trans, cis* + *trans*, and compensatory).

**Figure 4.**
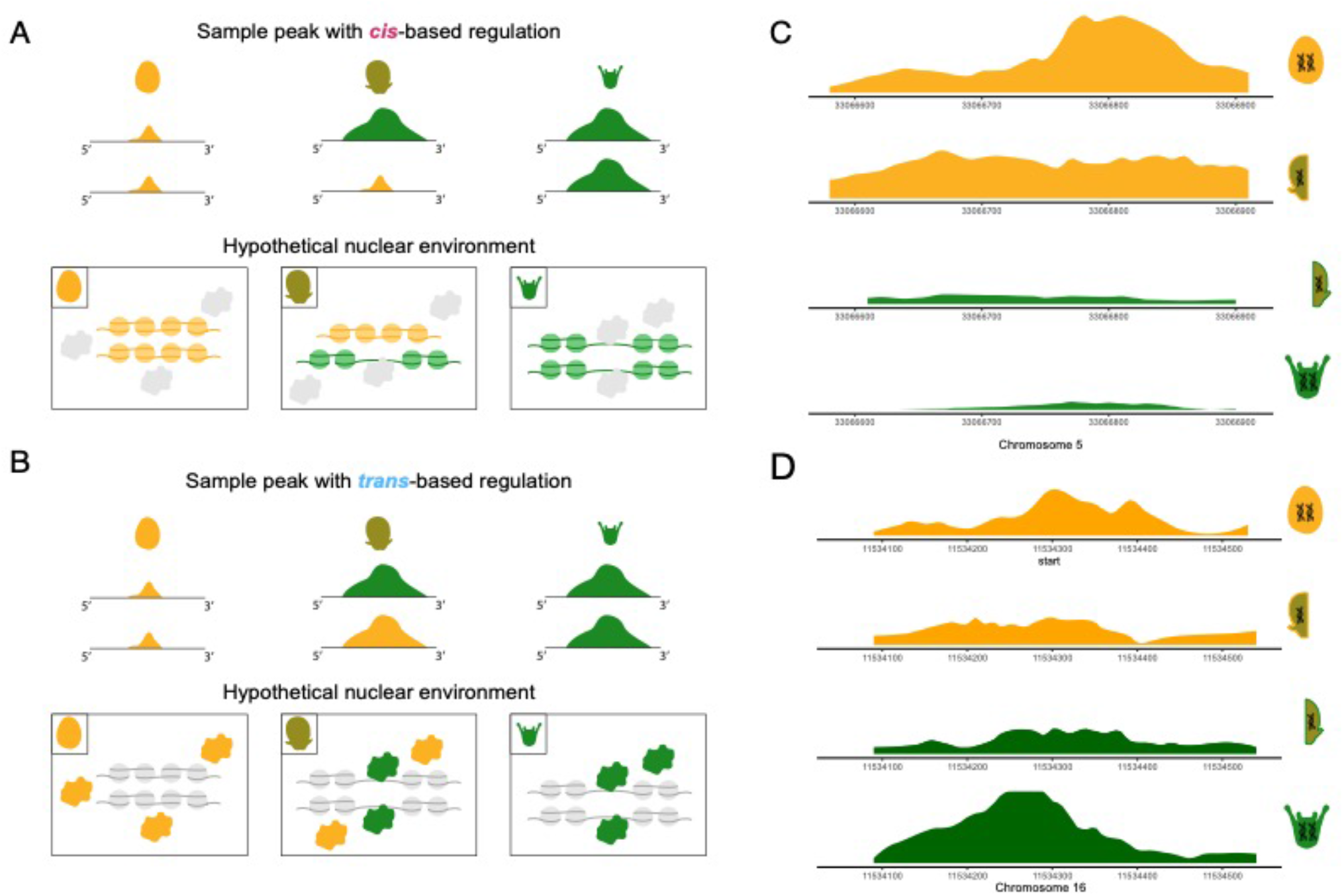
Models and examples of evolutionary change in regulatory mode. Left, theoretical examples of browser tracks for open chromatin (peaks) displaying *cis*- and *trans*-based regulation (A and B, respectively). For each cross, peaks on both maternal (orange) and paternal (green) alleles are shown. Directly below each is a model of the hypothetical interaction between *trans*-acting proteins, nucleosomes, and DNA that could potentially generate the pattern of chromatin accessibility in each cross. The color of the nucleosomes, DNA, and proteins indicate which species they were inherited from (gray indicates a lack of species-specific difference for that element). **(A)** Model of *cis*-based change. In the *He* same-species cross (left panels), DNA is tightly wound around nucleosomes, and thus *trans*-acting factors (proteins) are unable to interact with it, resulting in very small peaks on both alleles. In the *Ht* same-species cross (right panels), DNA is not wrapped around nucleosomes, leaving it accessible to *trans*-acting factors, and thus generating two large peaks in the corresponding browser tracks. In hybrids (center), one allele from each parent is inherited, yielding one large peak and one small peak as the two alleles differ in their ability to interact with trans-acting factors. **(B)** Model of *trans*-based change. In the *He* same-species cross (left panels), DNA is tightly wound around nucleosomes, and *trans*-acting factors are unable to interact with it, resulting in very small peaks on both alleles. In the *Ht* same-species cross (right panels), *trans*-acting factors (which differ from *He trans*-acting factors) are able to open up the chromatin and interact with it, thus generating two large peaks in the corresponding browser tracks. In the hybrid cross (center), *trans*-acting factors from each parent are present and able to interact with alleles inherited from either parent. Thus, the *Ht trans*-acting factors are able to open up the chromatin on both alleles, generating two equal-sized peaks in the browser tracks for the hybrid cross. Right, real examples of browser tracks corresponding to distinct regulatory modes. **(C)** *cis*-based change, **(D)** *trans*-based change. In both cases, total accessibility is shown for the same-species crosses, while for the hybrid, the browser track has been broken out into the accessibility of the two different alleles.

Most cases of differential accessibility between species are consistent with a single locus model (i.e., can be explained by a single mutation). Conversely, inferences that require multiple loci (e.g., additive, compensatory, and *cis* x *trans*) are relatively uncommon. These results hint at the possibility that the evolutionary divergence in chromatin states between the two *Heliocidaris* species often has a relatively simple genetic basis. Nonetheless, the variety of inheritance and regulatory modes inferred suggest that a variety of molecular mechanisms can contribute to evolutionary divergence in chromatin state during early development. Moreover, a substantial number of OCRs change inheritance or regulatory mode at least once across development (Figure S4), suggesting that the interplay between cis and trans effects is complex. The following paragraphs examine the variety of distinct genetic effects we observed, as well as and their relationship to the transcriptome.

### Maternal Dominance Effects Persist Longer in the Epigenome than the Transcriptome

With regard to inheritance mode, many more differential OCRs were classified as maternal dominant than paternal dominant (67,686 total maternal dominant peaks vs 17,568 total paternal dominant peaks). This ratio decreases slightly over developmental time (4.12, 3.80, 3.76 maternal:paternal at blastula, gastrula, larva). Early maternal dominance with a subsequent decrease during development is expected, as maternally provisioned regulatory molecules initially predominate in the nucleus but will be depleted over time as the zygotic genome (including paternal alleles) begins to exert an influence. Although we do observe this predicted decrease in maternal dominance effects in chromatin, the magnitude of the drop in accessibility is much less substantial than previously reported for the transcriptome across the same developmental stages: maternal dominance of mRNA abundance is overwhelmingly more common than paternal dominance at blastula, but maternal and paternal dominance are nearly equally represented by gastrula and even closer in larva (Figure 3A).

Maternal dominance effects on chromatin outnumber all other classifications of inheritance mode put together--even at larva, the latest developmental stage we examined. Indeed, the number of DA OCRs with maternal dominance is lowest in absolute terms at the earliest stage (blastula) and nearly doubles by the gastrula stage (15,580 to 27,381). Although it may seem paradoxical for maternal effects to increase over developmental time, this is made possible by the fact that more and more OCRs become accessible during development(Davidson, et al. 2022),resulting in the relative proportion of maternal dominant peaks remaining roughly stable over time despite an increase in their absolute number. These observations suggest that maternal effects on the chromatin landscape persist much longer during development than do maternal effects on transcript abundance, and are notably more extensive at later stages. This observation also indicates that dominance effects in the transcriptome do not directly parallel or reflect dominance effects in the chromatin landscape.

### Evolutionary Differences in the Epigenome are the Result of Extensive Cis and Trans Effects

Turning to regulatory mode, approximately equal numbers of differential OCRs are inferred to be all-*cis* and all-*trans* at the blastula stage (Figure 3B). The number of all-*cis* OCRs rises modestly at each subsequent stage of development, slightly outnumbering all-*trans* OCRs by larva stage. Unlike dominance effects, there is no clear *a priori* expectation about the proportion of *cis*- and *trans*-based contributions to evolutionary divergence in chromatin status, nor how these might change during development. That said, it is not surprising to find evidence of extensive *cis*- and *trans*-based genetic influences on individual OCRs, since a variety of changes in local sequence and in the *trans*-acting nuclear environment could in principle alter local chromatin configuration.

The overall trends in regulatory mode during development for the epigenome are generally different from, and in some cases the opposite of, trends in the transcriptome. First, the number of genes with an all-*cis* change in transcription progressively decreases with developmental time, whereas it increases for chromatin. Second, the difference in the number of all-*cis* and all-*trans* changes is greatest at blastula stage for transcription but lowest for chromatin (Figure 3B). Third, all-*trans* effects are nearly as common as all-*cis* effects in the transcriptome at blastula stage and then drop to nearly zero at gastrula and larva, but in chromatin all-*trans* effects remain fairly constant across all developmental stages. Finally, the three regulatory mode classifications that imply a genetic influence from multiple loci (*cis* + *trans*, *cis* x *trans*, and compensatory) are generally less common for chromatin than for the transcriptome. In particular, *cis* x *trans* effects are moderately prevalent in the transcriptome at all three stages, but consistently rare for chromatin. Most dramatically, the extensive compensatory effects seen in the blastula stage in the transcriptome—which are likely due to maternal effects—are completely absent from chromatin. Alone among the multi-locus inferences, *cis* + *trans* effects are similar in the transcriptome and chromatin, where they are rare at all stages examined. Together, these observations indicate that the inferred regulatory modes underlying evolutionary changes in the transcriptome do not parallel those in the chromatin landscape.

### Large differences in chromatin accessibility are often based in cis

We next considered the magnitude of genetic effects on chromatin accessibility. Specifically, we compared *cis*- and *trans*-based influences on accessibility of OCRs. We found that, at all stages, peaks with *cis*-based differences in accessibility had a greater (absolute) difference in accessibility between species than did peaks with *trans*-based differences in accessibility (Figure S5A, p<<0.05 at all three stages). Moreover, when peaks were ordered by the magnitude of these effect sizes, an average of 63% of the top 10 peaks were *cis* (Figure S5B,C). These results indicate that local mutations are as, if not more, important for evolutionary changes in local chromatin landscape relative to *trans* effects.

### Genetic basis of differential chromatin predicts differential expression

Next, we turned our attention to understanding how evolutionary changes in the epigenome influence the transcription of nearby genes. Specifically, we asked whether differential chromatin accessibility is enriched near differentially expressed genes and vice-versa. (Note that these are not symmetrical tests, due to the many-to-one relationship between regulatory elements and genes.) First, we considered a “peaks focused” perspective: if a peak is differentially accessible, is the single nearest gene more likely to be differentially expressed? We found that, at all three stages, differentially accessible peaks had a nearest gene that was differentially expressed more often than expected by chance (Chi-squared test for independence) (Figure 5A, S6A). Second, we considered a “gene-focused” perspective: if a gene is differentially expressed, is there any “nearby” (within 25 kb) chromatin peak that is differentially accessible? At the blastula stage, there was not a statistically significant association between differentially expressed genes and nearby differentially accessible peaks from either the “peaks-focused” or “genes-focused” perspective. However, at the later two stages we did observe that differentially expressed genes had at least one nearby differentially accessible peak more often than expected by chance (Chi-squared test for independence, Figure 5B, S6B). Moreover, the strength of these correlations increased during development for both the “peak focused” and “gene focused” comparisons, and were notably strongest at larva, when the zygotic genome is the most extensively transcribed and when maternal effects are the weakest among the stages examined.

**Figure 5.**
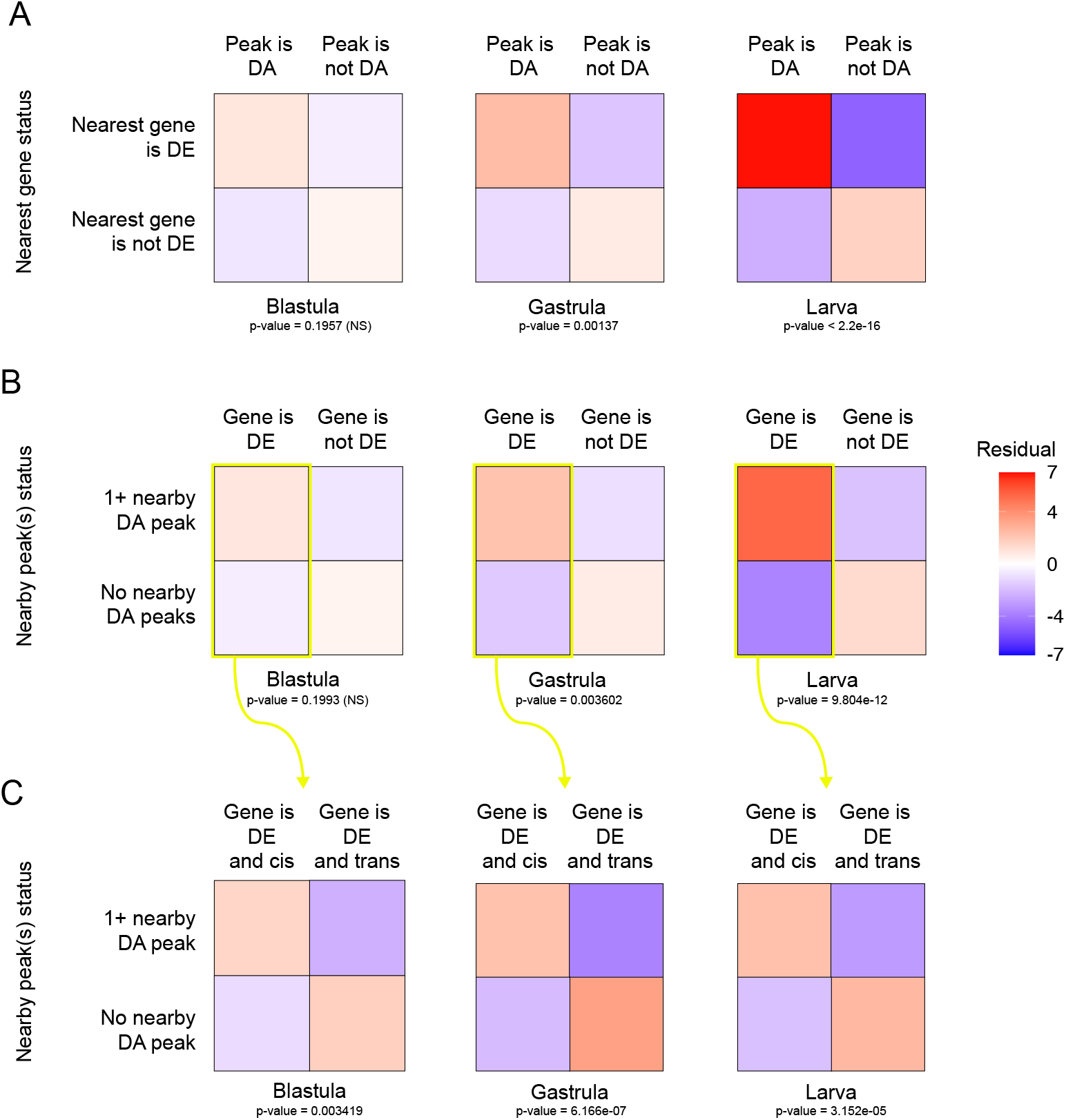
Relationship between evolutionary change in chromatin configuration and transcript abundance. Chi-squared tests for independence were used to measure the correlation between evolutionary changes in open chromatin regions and in expression of nearby genes. Heatmaps show residuals for tests carried out in three different contexts (see Supplementary Figure 4 for details). Larger residual values (darker red squares) indicate an enrichment and suggest that there are more of the given event than expected by chance. Tests were carried out separately at each of the three developmental stages. **(A)** The “peaks-focused” tests ask whether the nearest gene to a differentially accessible peak is itself differentially expressed more often than expected by chance. The chi-squared tests were significant (test statistic p<0.05) at gastrula and larva. **(B)** The “gene-focused” tests ask whether there is at least one differentially accessible peak within 25 kb of a differentially expressed gene more often than expected by chance. The chi-squared tests were significant (test statistic p<0.05) at gastrula and larva. Note that “peaks-focused” and “gene-focused” tests are not redundant, due to the 1-to-many relationship between genes and regulatory elements. **(C)** The “regulatory mode-focused” tests were carried out for genes that are differentially expressed between species. These tests ask whether *cis*- and/or trans-based differential expression of genes are enriched for differentially accessible peaks within 25 kb more often than expected by chance. The chi-squared tests were significant (test statistic p<0.05) at all three stages, with *cis*-based differential expression enriched for nearby DA peaks and *trans*-based differential expression depleted for nearby DA peaks.

We further interrogated the connection between differentially-expressed genes and differentially accessible chromatin by asking whether *cis*- or *trans*-based DE genes were more associated with nearby differentially accessible chromatin. We reasoned that *cis*-based differential expression *should* be enriched near differentially accessible chromatin (since this is one possible molecular mechanism contributing to evolutionary changes in transcription), while *trans*-based differential expression should not be enriched (since local genetic differences are inferred to have little to no contribution to evolutionary changes in transcription). We therefore examined OCRs near genes inferred to have *cis*-based variation in gene expression, in order to assess how much of the variation in the expression of these genes can be attributed to changes in the accessibility of nearby potential *cis*-regulatory elements. We found that genes with *cis*-based differences in expression were enriched for at least one differentially accessible nearby OCR at all three stages examined (Chi-squared test for independence). Meanwhile, genes with *trans*-based differences in expression did not show any such enrichment (Figure 5C, S6C).

Together, these results suggest that some evolutionary changes in the chromatin landscape contribute to evolutionary changes in transcription during development in *Heliocidaris*. The enrichment of *cis*-based differences in chromatin accessibility near differentially expressed genes further suggests that some of this influence is genetically based.

### Proximal and distal peaks differ in size, regulatory mode, and motif enrichment

Given that gene regulatory elements carry out diverse functions, we investigated whether OCRs show distinct evolutionary properties based on their function as core promoters and distal enhancers. We used position relative to the nearest TLS as a proxy for likely function, dividing OCRs into proximal peaks (center ≤500 bp from a TLS) and distal peaks (center >500 and <25000 base pairs from a TLS). In total, there were 3,474 proximal peaks and 88,560 distal peaks. Both proximal peaks and distal peaks had significantly higher rates of differentially accessible peaks than other peaks in the peak set at all three stages (Fisher’s exact test, Table S8). When we examined the regulatory modes of these differentially accessible peaks, we found that proximal peaks are more than twice as likely to be genetically based in *trans* than distal peaks, and are also enriched for *trans* peaks relative to the proportion of *trans* peaks in the peak set as a whole (Chi-squared test for independence and Fisher’s exact test, Fig 6A, Table S9). Proximal peaks also had a significantly greater effect size than distal peaks at all stages studied (Welch’s t-test, Fig 6B), where effect size is defined here as the log2 of the ratio between the accessibility of the same-species peaks (as in (Mattioli, et al. 2020; Wang, et al. 2020)).

**Figure 6.**
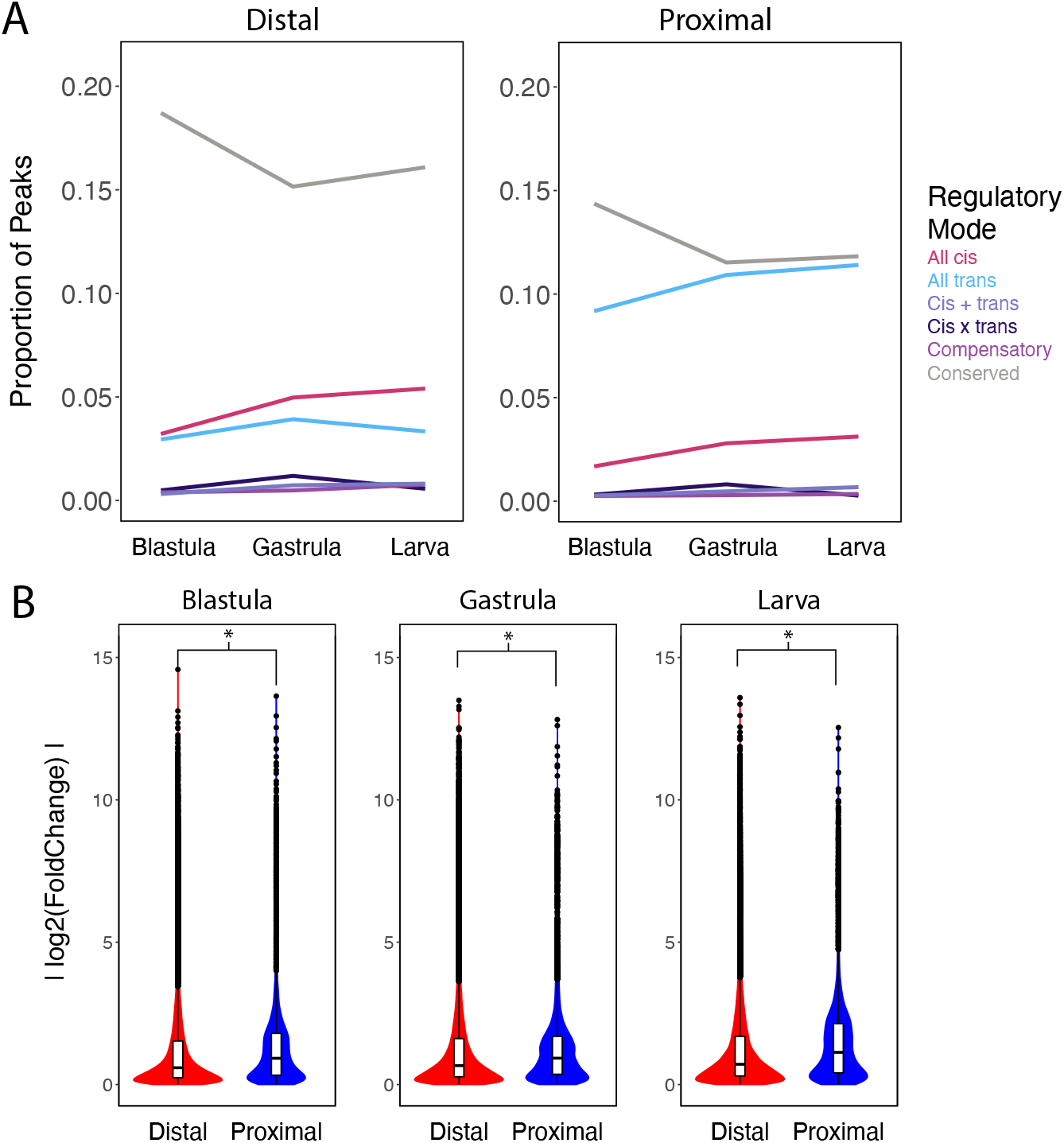
Distinct evolutionary trends in proximal and distal open chromatin regions. (A) Line plots showing the proportion of open chromatin regions in each regulatory mode classification for proximal (<500 bp from the translation start site (TLS) of the nearest gene) *vs* distal (between 500 bp and 25 kb from TLS). *Trans*-based differences dominate proximal peaks, while *cis*-based differences are slightly more common in the much more abundant distal peaks. (B) Violin plots contrasting the effect size for distal *vs* proximal peaks. At each stage, the mean effect size for proximal open chromatin regions was significantly greater than the mean effect size for distal open chromatin regions (Welch’s t-test, blastula: p=4.485e-11; gastrula: p=0.01414; larva: p<2.2e-16).

To assess the molecular mechanisms controlling access to proximal and distal elements, we carried out motif enrichment using the HOMER motif analysis tool. Based on our previous analysis (Davidson, et al. 2022), we expected to see enrichments for motifs related to pioneer factors in at least the proximal peak subset. We found that nearly 800 motifs were enriched in proximal elements as compared to distal elements, while only 8 motifs were enriched when distal elements were the test set (Table S10). Among the motifs enriched in proximal elements were those for several Forkhead family transcription factors, including FOXK1, FOXO3, and FOXF1.

### Peaks near gene-regulatory network (GRN) genes have a unique signature

Transcriptional states in sea urchin embryos are driven by a well-defined gene regulatory network (Davidson, et al. 2002; Oliveri, et al. 2002; Saudemont, et al. 2010; Erkenbrack, et al. 2018). Previous work has established that 1) the genetic mechanisms governing the regulation of these genes differ from those controlling gene expression as a whole (Wang, et al. 2020), and 2) putative regulatory elements near these genes are under greater selective pressure than the rest of the epigenomic landscape (Davidson, et al. 2022).Thus, we also examined how the open chromatin landscape near GRN genes compared to the epigenome as a whole. We found that the density distribution of peaks near (within 25 kb) of a GRN gene was significantly different from the density distribution of peaks in the entire genome (Scheffe test and Kolmogorov-Smirnov test), with GRN genes having more nearby peaks than non-GRN genes. While this difference in the epigenomic landscape may be due to a unique property of GRN genes themselves, it is also possible that it is a consequence of the fact that the vast majority of GRN genes are transcription factors. To test this possibility, we compared the density distribution of peaks near GRN genes to that of peaks near all transcription factors. We found that the GRN genes did not tend to have a greater number of nearby peaks (Fig 7A) than the entire set of transcription factors.

**Figure 7.**
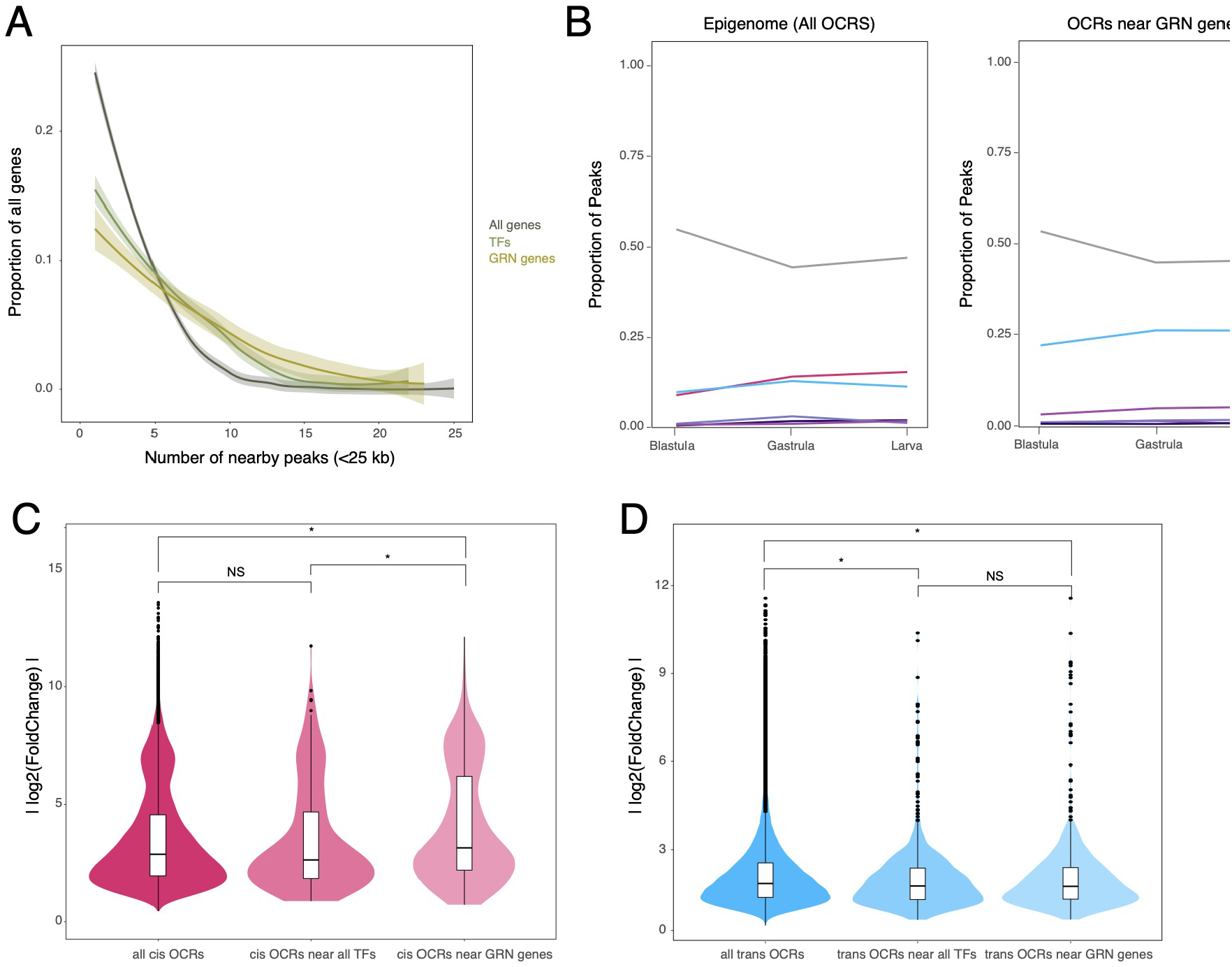
Distinct evolutionary trends in open chromatin near developmental regulatory genes. Plots present results for genes and open chromatin regions in three classes of interest (“all genes”, “transcription factors” and “GRN genes”). **(A)** Smoothed histograms of the proportion of genes with a given number of nearby peaks. The distributions for “transcription factors” and “GRN genes” were both significantly different from the distribution for “all genes” by a Kolmogorov-Smirnov test (p=1.179e-04 and p=3.108e-05 respectively), but not significantly different from each other. The X axis was truncated to 25 for illustration purposes (all 3 distributions are heavily right-skewed, with a tiny proportion of values >25). **(B)** Line plots of regulatory mode classification for all peaks (left) as a proportion of the total number of peaks *vs* for peaks within 25kb of a GRN gene (right). **(C)** Violin plots of effect size for *cis*-based peaks in three classes of interest. Mean effect size for *cis*-based peaks near GRN genes was significantly greater than the mean effect size for *cis*-based peaks near any transcription factor (Scheffe test, p=1.0e-04), and also significantly greater than the mean effect size for *cis*-based peaks in general (Welch’s t-test, p=3.304e-07 and Scheffe test, p=1.8e-10). The mean effect sizes of the latter two categories did not significantly differ from each other. **(D)** Violin plots as in (C) but for *trans*-based peaks in the same three classes of interest. Here, the mean effect size for *trans*-based peaks near GRN genes was significantly smaller than the mean effect size for *trans*-based peaks in general (Scheffe test, p=2.4e-9), but was not significantly different from the mean effect size for *trans*-based peaks near any transcription factor. The mean effect size for trans-based peaks near any transcription factor was also significantly smaller than for trans-based peaks in general (Scheffe test, p=0.00022).

To provide a richer insight into the connection between the regulation of GRN genes and the regulation of nearby regulatory elements themselves, we next queried the regulatory modes of peaks near GRN genes. These peaks were, at all three stages, approximately twice as likely to be *trans* compared to the epigenome as a whole (Fisher’s exact test, Figure 7B). They were also more likely to be *trans* than peaks near transcription factors as a whole (Fisher’s exact test). As with the whole epigenome, *cis* peaks had a significantly greater effect size than *trans* peaks at all stages studied (Welch’s t-test). However, *cis* peaks near GRN genes were also more open than *cis* peaks as a whole (as well as *cis* peaks near transcription factors), while *trans* peaks near GRN genes were less open than *trans* peaks as a whole (Scheffe test and Welch’s t-test, Figure 7C and D). Thus, peaks near GRN genes are both more numerous than near comparable genes (i.e., those encoding transcription factors not part of the GRN), and also more likely to have a *trans*-based difference in accessibility between species. At the same time, the magnitude of the difference in accessibility between species is markedly smaller for *trans* peaks than *cis* peaks, a contrast which is also true at the level of the entire epigenome but which is exaggerated when only peaks near GRN genes are considered.

Similarly to our analysis of proximal vs. distal peaks, we compared motif enrichment in peaks that were *cis* in at least one stage and peaks that were *trans* in at least one stage to probe what molecular mechanisms might be controlling the regulation of these peaks. *Cis* peaks were enriched in 116 motifs when *trans* peaks were used as the background set. Of these motifs, 6 matched known GRN genes. When *cis* peaks were used as the background set, *trans* peaks were enriched in 301 motif, with 32 of these motifs matching known GRN genes. (Table S11).

## DISCUSSION

We present here the first analysis of chromatin configuration in interspecies hybrids using outbred natural populations and covering multiple stages of embryonic development. An important goal of this study was to understand the potential for evolutionary changes in DNA accessibility to influence trait evolution by modifying gene expression. While it is mechanistically plausible that changes in the accessibility of regulatory elements contribute to trait differences between species, there is little published evidence. For this to be true, three minimal conditions must hold: (1) chromatin status differs consistently between species, (2) those differences influence gene expression, and (3) they are genetically based. In prior work with the same species and developmental stages, we found that thousands of OCRs differ in accessibility between species and that these changes are concentrated in OCRs near genes encoding transcription factors and specifically on the branch leading to the derived life history (Davidson et al. 2022a, 2022b). These studies also found a correlation between divergence in chromatin status and divergence in the expression of nearby genes. Together, these results address the first and second conditions, and further suggest that changes in chromatin accessibility contributed to the life history shift within *Heliocidaris*. In the present study, we extend evidence in support of the second condition and, for the first time, investigate the critical third condition, namely the genetic basis for evolutionary changes in chromatin accessibility and their influence on gene expression during development. Our findings can be summarized by five major themes, which we highlight below.

### Many differences in chromatin accessibility are genetically based

Chromatin configuration can differ substantially across life history stages, cell types, and environmental conditions even within a single genotype. This raises an important question in an evolutionary context: are differences in chromatin observed between closely related species genetically based, or do they simply reflect plasticity in response to an altered nuclear environment? This question matters for understanding how natural selection operates on chromatin status, because the more direct the genetic basis is for a trait, the more readily natural selection can act on that trait. The issue is particularly acute for *Heliocidaris*, as eggs of the two species differ enormously in the transcripts, proteins, and metabolites that are loaded into the egg (Hoegh-Guldberg and Emlet 1997; Byrne, et al. 1999; Israel, et al. 2016; Davidson, et al. 2019). For this reason, it is plausible that differences in chromatin status between the two species arise largely due to indirect effects arising from different nuclear environments, rather than arising from the genetics of the developing embryo. Plasticity of chromatin accessibility in response to differing nuclear environments is likely to occur via changes to the expression or localization of transcription factors that interact with chromatin—in other words, due to *trans* changes (Figure 1). On the other hand, differences in chromatin accessibility that are based in *cis* must be genetic. By placing chromosomes in the common nuclear environment of hybrid embryos, we were able to measure the relative contributions of *cis* versus *trans* changes to chromatin accessibility, and we found that slightly over half of the differences in open chromatin between species in *Heliocidaris* have a *cis* effect (Figure 3B). Thus, a substantial number of the observed differences in chromatin accessibility between our two species is based in genetics and can be acted upon via evolutionary mechanisms. Moreover, *cis* changes by definition are local to each instance of differential chromatin—unlike *trans* changes, they cannot be explained by a change to a single upstream regulatory factor. Therefore, the widespread nature of *cis* changes in the epigenome suggests that evolutionary modifications to chromatin accessibility occurred through numerous local mutations rather than one or a few a modifications in upstream factors.

### Changes in chromatin accessibility are associated with changes in gene expression

When considering the contribution of evolutionary changes in chromatin configuration to the evolution of gene expression, two important caveats should be kept in mind. First, a difference in chromatin status does not by itself indicate an influence on transcription. From a mechanistic perspective, opening chromatin is permissive rather than determinative: unless the appropriate transcription factors are present, a change in accessibility alone will not alter transcription. This is clearly illustrated by the observation that many OCRs open prior to the onset of transcription of any nearby gene during development, including in sea urchins specifically (Shashikant, et al. 2018b; Davidson, et al. 2022). Second, a change in chromatin configuration is only one of several molecular mechanisms that could alter gene expression: other possibilities include a change in the sequence of the gene itself and in a variety of post-transcriptional processes

Despite these caveats, we found a statistical association between differential chromatin status and differential gene expression at two of the three stages of development examined. The strength of this association increased over developmental time, likely reflecting the maternal-to-zygotic transition in the mRNA pool and the progressive appearance of zygotically synthesized transcription factors during development. As expected, the statistical association is stronger for genes whose genetic basis for expression is in *cis* than for those whose genetic basis is in *trans* (Figure 5C, Figure S6). Together, these results suggest that evolutionary changes in chromatin configuration contribute to some evolutionary changes in gene expression; moreover, the fact that we could detect such correlation at all, given the (Shashikant, et al. 2018b)caveats just laid out, suggests that the role of chromatin configuration changes in changes to gene expression is not insubstantial and may in fact be one of the primary drivers of changes in gene expression.

### Genetic mechanisms controlling changes in chromatin status show different patterns from those controlling gene expression

We found that the regulatory modes governing changes in chromatin status are markedly different from those controlling changes in gene expression. The number of genes with no differential expression (genes with a “conserved” regulatory mode) increased through development, indicating that the transcriptomes of the two species appear to converge as the embryos reach metamorphosis. However, the epigenome maintains and actually increases species-specific differences in accessibility as development progresses, as indicated by the fact that sites with a “conserved” accessibility status decrease from blastula to larva. Thus, it would appear that many differences in chromatin accessibility do not feed forward into changes in the expression of nearby genes; indeed, other studies (Connelly, et al. 2014) suggest that this may be the case. On the other hand, it does appear that there is at least a loose relationship between differential accessibility and differential expression, as differentially accessible peaks are enriched near differentially accessible genes and vice versa. Moreover, genes with *cis*-but not *trans*-based differences in expression are enriched for nearby differentially accessible chromatin, suggesting that that the mechanism driving sequence-based differences in expression may be located in nearby enhancer elements. This finding also provides evidence that knowing the inferred genetic basis behind a difference in gene expression can help strengthen the ability to discover instances of differential chromatin accessibility that may be mediating the difference in gene expression.

It is also possible that these species-specific differences in accessibility do have functional relevance, but only for later stages beyond the time course studied here. Such a result would not be without precedent, as previous work has shown that the epigenome often becomes accessible hours before associated genes are activated (Shashikant, et al. 2018a). Overall, it appears that differentially accessible chromatin is permissive of but not always causal to changes in gene expression.

Dominance patterns also persist in the epigenome longer than in the transcriptome, as evidenced by the fact that the number of maternally and paternally dominant genes converges as development progresses, while the difference between the number of maternally and paternally dominant peaks remains fairly static over time. This indicates that a strong maternal influence on the epigenome lingers even after the maternal-to-zygotic transition has largely eroded the effects of maternal deposition in the transcriptome. Thus, this pattern also suggests that changes in chromatin status can evolve independently from changes in gene expression. The discordance between inheritance patterns in the transcriptome and the epigenome may be another example of epigenomic changes being facilitative of, but not necessarily directly causal to, changes in gene expression, as mentioned earlier. Moreover, these persistent maternal effects at the level of the epigenome are consistent with findings in our previous work (Davidson, et al. 2022) that suggest fewer active changes in chromatin status across development in *H. erythrogramma* relative to *H. tuberculata*. While more work must be done to fully understand the meaning of these results, this analysis nevertheless underscores the importance of investigating the inheritance of chromatin accessibility, as these results were not predictable based on previous work.

### Cis peaks have larger between-species differences in accessibility than trans peaks

When different regulatory modes were compared, we found that peaks with regulation based in *cis* had a greater average difference in accessibility between same-species crosses than did peaks with regulation based in *trans*. This finding is similar to those seen in yeast hybrids (Ronald and Akey 2007; Connelly, et al. 2014) but is, to our knowledge, the first time this observation has been documented among species in wildtype multicellular eukaryotes. Given that *cis* changes can evolve quickly and have large influences on enhancer and promoter activity (Yona, et al. 2018; Kurafeiski, et al. 2019), we propose a model in which *cis*-based changes in general cause large increases or decreases in accessibility at a single site, while *trans* changes, which are more likely to be pleiotropic (Carroll 2005; Chesler, et al. 2005) and cause smaller changes in the accessibility of any individual OCR.

### Cis changes implicated in most striking changes to chromatin accessibility between species

We demonstrate that while both *cis* and *trans* influences have extensive effects on species-specific differences in chromatin accessibility, *cis* changes appear to exert a larger influence on the chromatin landscape overall, indicating that there is a strong genetic basis for differential chromatin accessibility. This conclusion is supported by the fact that *cis* effects occur with slightly greater frequency than *trans* effects at all three of the developmental stages studied, have larger average effect sizes than *trans* effects, and are overrepresented in the set of the most differentially accessible regions of the genome. Moreover, while paternally dominant changes in accessibility are rare, they are measurable and increase with developmental time, indicating that the paternal genome can and does influence chromatin accessibility during development. Combined, these results suggest that differences in chromatin accessibility between species are not due entirely, or even mostly, to differences in maternal provisioning. Nevertheless, there remains a striking role for *trans* factors in the accessibility of biologically relevant peaks, as described below.

### Chromatin near GRN genes differs from the rest of the epigenome in important ways

We considered how the accessibility of peaks near GRN genes was regulated, and found marked differences in regulation patterns for these peaks versus the epigenome as a whole. This was manifested in at least three different ways: first, GRN genes were more likely than the transcriptome overall to have nearby differentially accessible regions; second, these regions were more likely to be *trans* than the epigenome as a whole; third, despite this, the difference in accessibility across species for *cis* vs. *trans* peaks near GRN genes was greatly exaggerated compared to this difference when *cis* and *trans* regions of the entire epigenome were compared. We submit that this observation is due to the level of importance of the GRN relative to the rest of the genome (Halfon 2017), leading to an exacerbated difference in accessibility between *cis* and *trans* peaks near GRN genes vs. *cis* and *trans* peaks in the rest of the epigenome. Furthermore, we would expect that peaks near GRN genes would be quickly selected for or against depending on the net advantage or disadvantage they create for the organism. This would lead to the observations that *cis*-based accessibility differences near GRN genes are relatively rare, but large in magnitude where they do occur. We also noted that when motif enrichments in *cis* and *trans* peaks were compared, the set of *trans* peaks was enriched for GRN motifs relative to the set of *cis* peaks. This would suggest that peaks which are regulated in *trans* may be more likely to be influenced by changing aspects of the GRN (which are themselves often transcription factors) than are peaks regulated in *cis*.

### Concluding thoughts

In this study, we examine how *cis* and *trans* factors contribute to patterns of chromatin accessibility in the developing embryo of two sea urchin species with markedly different life history strategies, and compare these findings to the genetic mechanisms governing gene expression during the same period of development. We find that, though differential chromatin accessibility is predictive of differential gene expression, particularly for genes with *cis*-based changes in expression, the transcriptome and epigenome are regulated very differently throughout development. *Cis* and *trans* factors both have striking effects on accessibility patterns, with *cis*-based effects being generally larger in magnitude and scattered throughout the epigenome, whereas *trans* factors are relatively smaller and disproportionately influence chromatin near genes involved in the dGRN. Interestingly, these regions of the genome whose accessibility is governed by *trans* factors also show evidence of enrichment for sequence-based motifs related to the dGRN. Together, these findings illustrate that understanding the genetics of the mechanisms regulating the epigenome can in turn further our understanding of the process that fine-tunes the regulation of gene expression. Moreover, they provide the first application of regulatory and inheritance mode interrogation in the epigenome of an outbred wild species pair with differing life history modes, and finally, they demonstrate the complexity of the interplay between *cis* and *trans* acting factors and the roles they play in regulating chromatin status.

## DATA AND RESOURCE AVAILABILITY

FASTQ Files of raw ATAC-seq reads are available on NCBI’s Sequence Reads Archive (accession number TBD). Count tables, lists of relevant gene sets for testing, and R code used to generate figures and tables are available on Dryad at (accession number TBD).

## ACKNOWLEDGEMENTS

We are grateful to the staff of the Sydney Institute of Marine Sciences for providing laboratory space, resources for experiments, and animal care. Krista Pipho, Micah Dailey, Alejo Berrio, Christina Zakas, and Nathan Harry provided feedback on manuscript drafts and figures. We also thank the staff of the Duke Sequencing and Genomic Technologies Shared Resource for assistance with sequencing. This work was supported by Training Grant T32GM007184-43 from the National Institutes of Health and a Graduate Research Fellowship from the National Science Foundation to HRD and by a grant from the National Science Foundation’s Division of Integrative Organismal Systems (award no. 1929934) to GAW.

## AUTHOR CONTRIBUTIONS

HRD and GAW conceived and designed the study. HRD, PLD, and MB performed sample collection. HRD performed the data analysis. HRD and GAW wrote the manuscript. All authors revised the manuscript.

