## Supplemental Figures for "Hybrid epigenomes reveal extensive local genetic changes to chromatin accessibility contribute to divergence in embryonic gene expression between species"

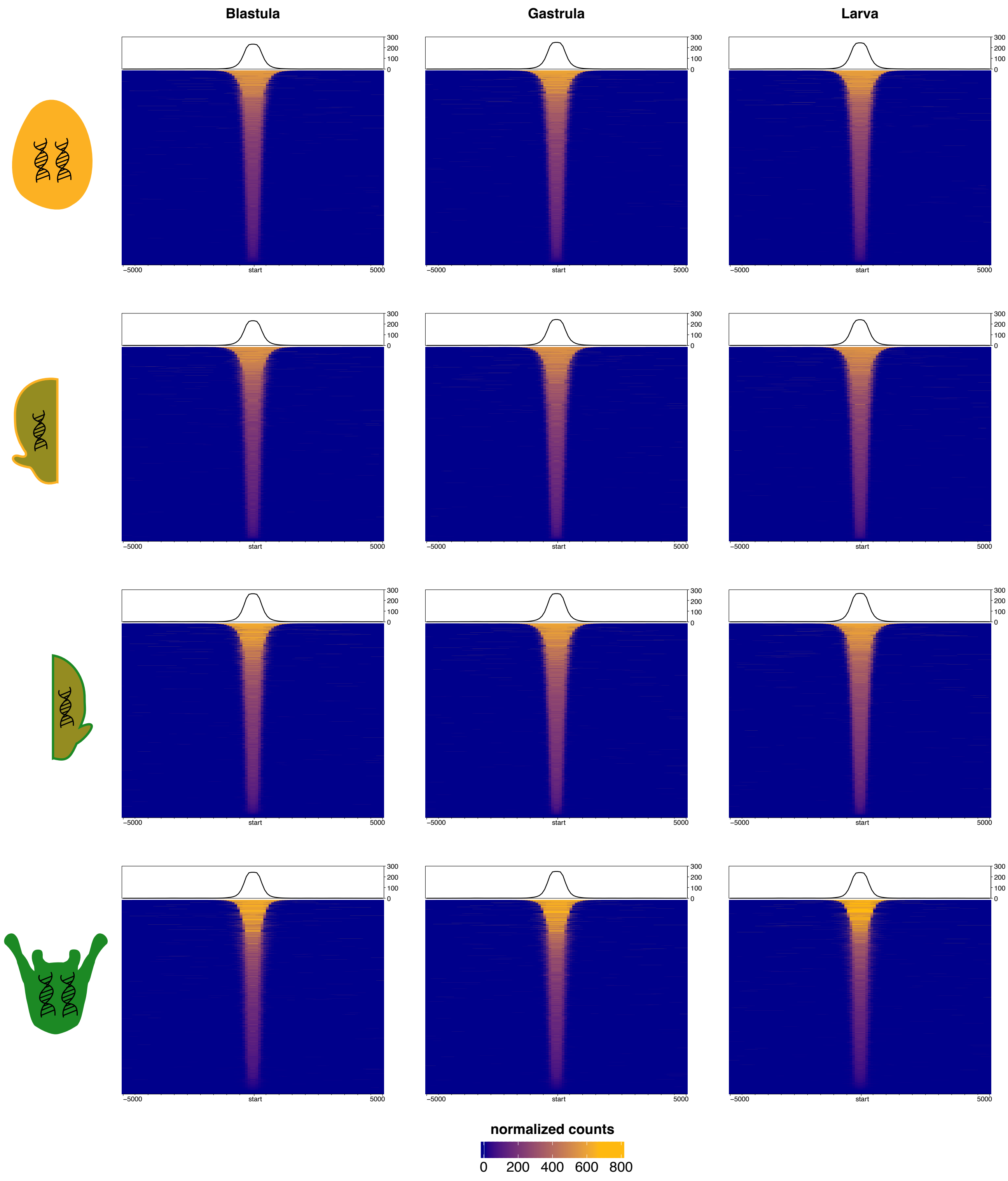


**Supplementary Figure 1.** Plots of peak size distribution for peaks <501 bp from a TLS, made using the "enrichedHeatmap" package in R.

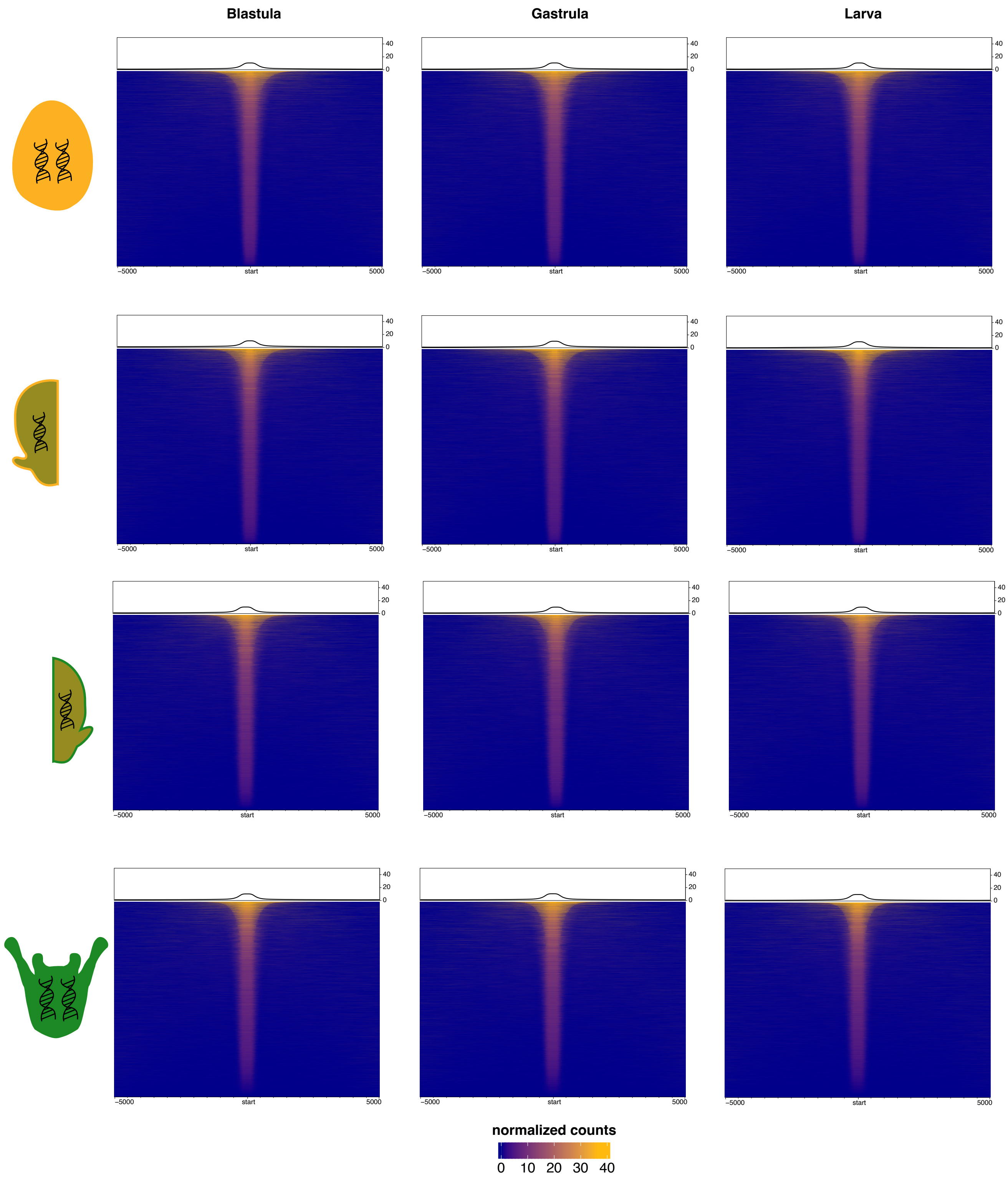


**Supplementary Figure 2.** Plots of peak size distribution for peaks between 501 and 25000 bp from a TLS, made using the “enrichedHeatmap” package in R. Note that the color scale for this figure differs from that of Supplementary Figure 1.

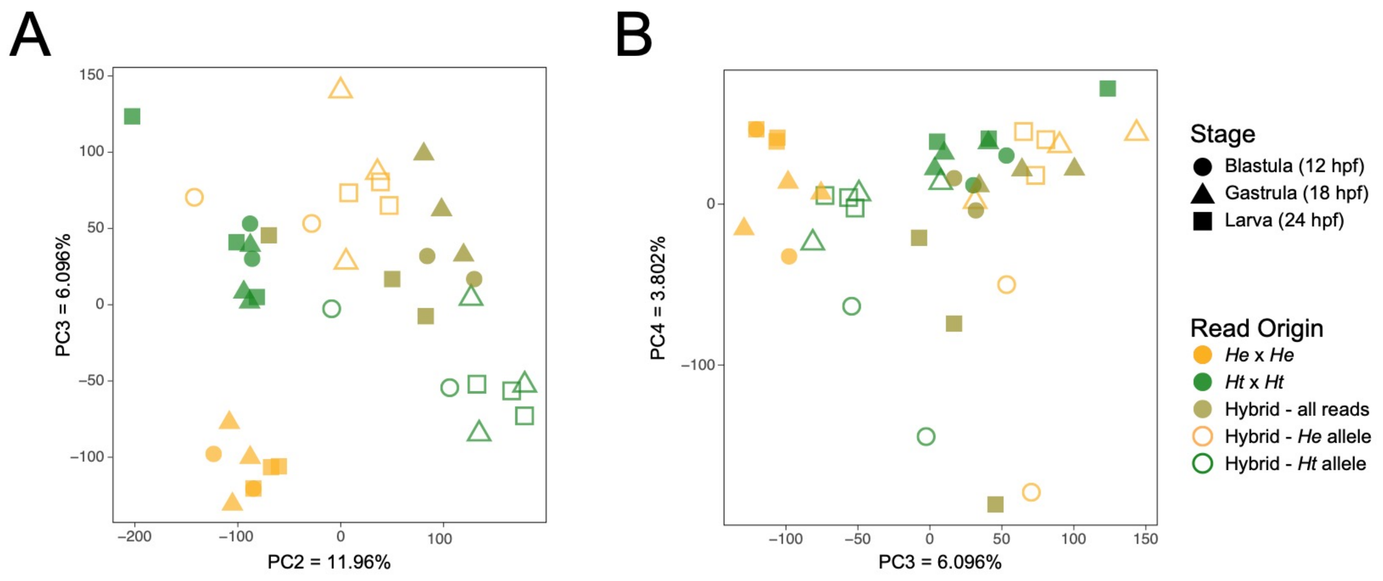


**Supplementary Figure 3.** Additional principal components (PCs) do not separate ATAC-seq reads by stage. **A)** PCA of PC2 vs PC3. **B)** PCA of PC3 vs. PC4. (For PC1 vs PC2, see Figure 2).

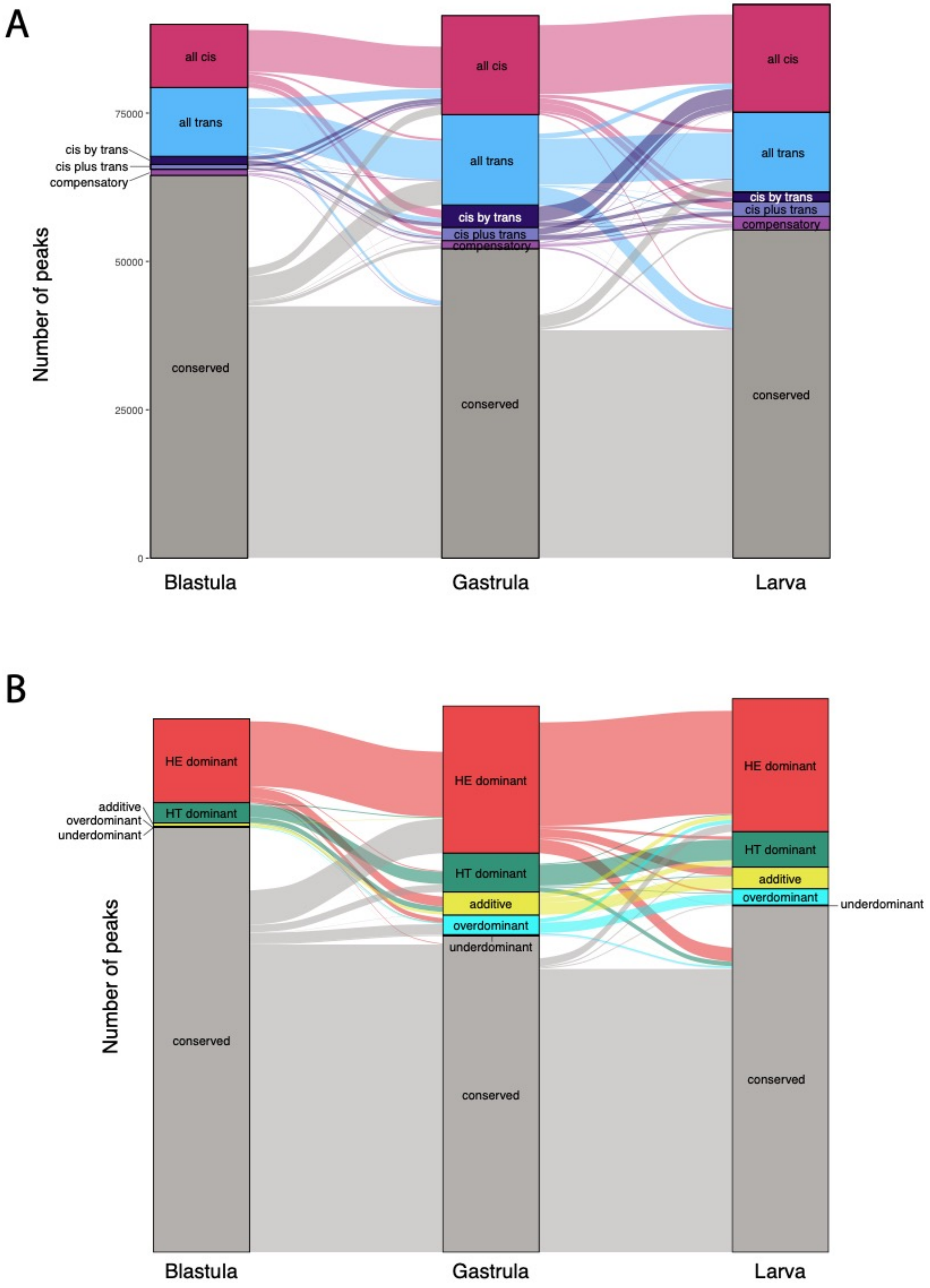


**Supplementary Figure 4.** Alluvial plots showing in the genetic basis for evolutionary differences in peaks across developmental time. **A)** Regulatory mode. **B)** Inheritance mode. Note that, for clarity, ambiguous peaks which differ by stage are not displayed; therefore, peak totals from one stage to the next are not equivalent.

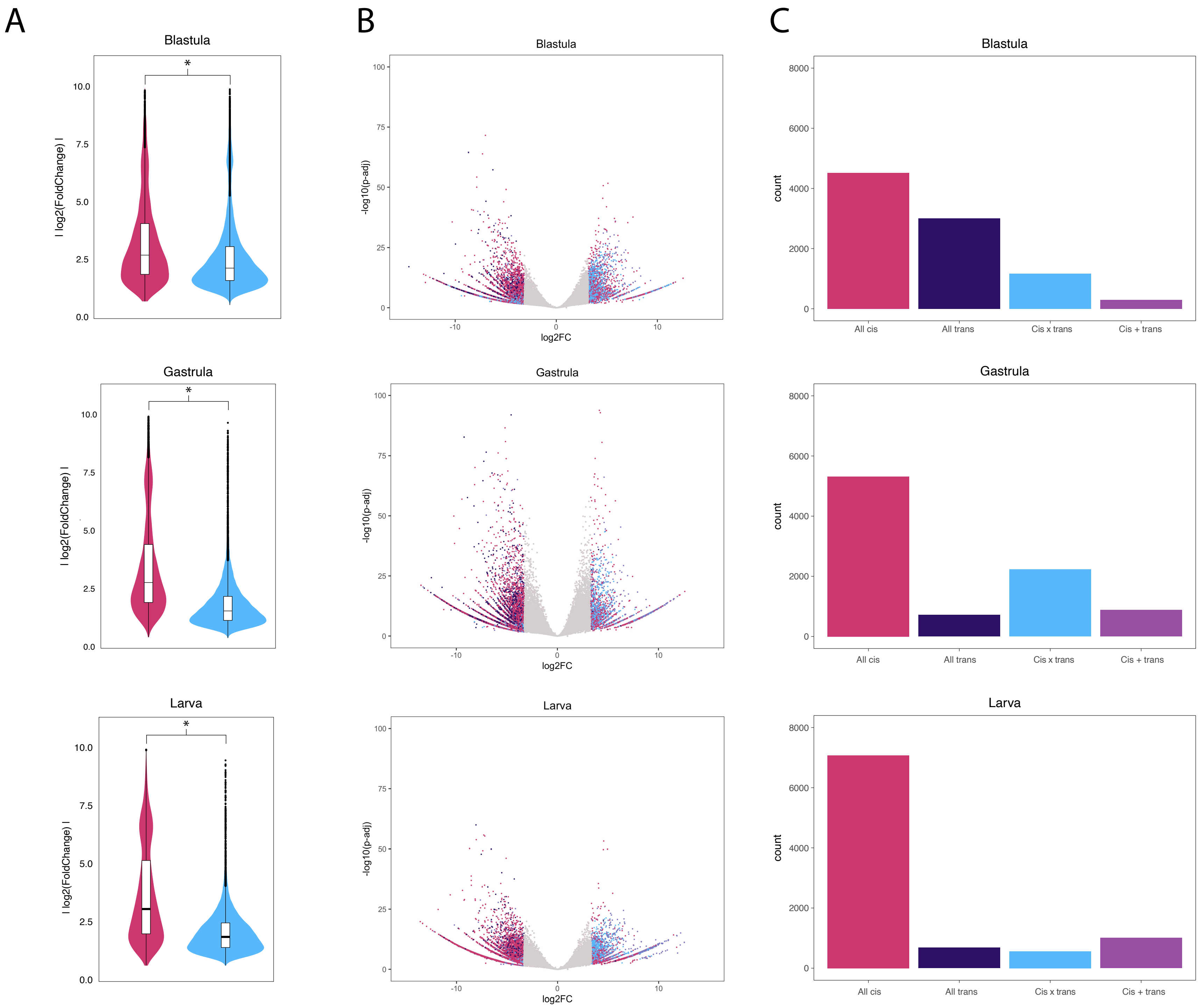
A
B
C
Blastula
Gastrula
Larva
I log2(FoldChange) I
*
-log10(p-adj)
log2FC
count
All cis
All trans
Cis x trans
Cis + trans

**Supplementary Figure 5.** Effect sizes in *cis*- and *trans*-based differential open chromatin regions. (**A)** Violin plots of the effect size (measured as the absolute value of the log2 fold change between the same-species crosses) for *cis*- *vs trans*-based open chromatin regions at the three developmental stages examined. At each stage, the mean effect size for *cis*-based open chromatin regions was significantly greater than the mean effect size for *trans*-based open chromatin regions (Welch's t-test, $p<2.2e-16$ for all three stages). **(B)** Volcano plots of open chromatin regions at each stage, stratified into the top 10% of effect sizes and the bottom 90% of effect sizes. Peaks in the top 10% for effect size are colored by regulatory classification, while peaks in the bottom 90% are light grey. (Compensatory and conserved peaks are not shown by color, as they fall within the bottom 90% of effect sizes by definition.) **(C)** Bar chart of the regulatory classification for peaks in the top 10% of effect size, colored as in (B).

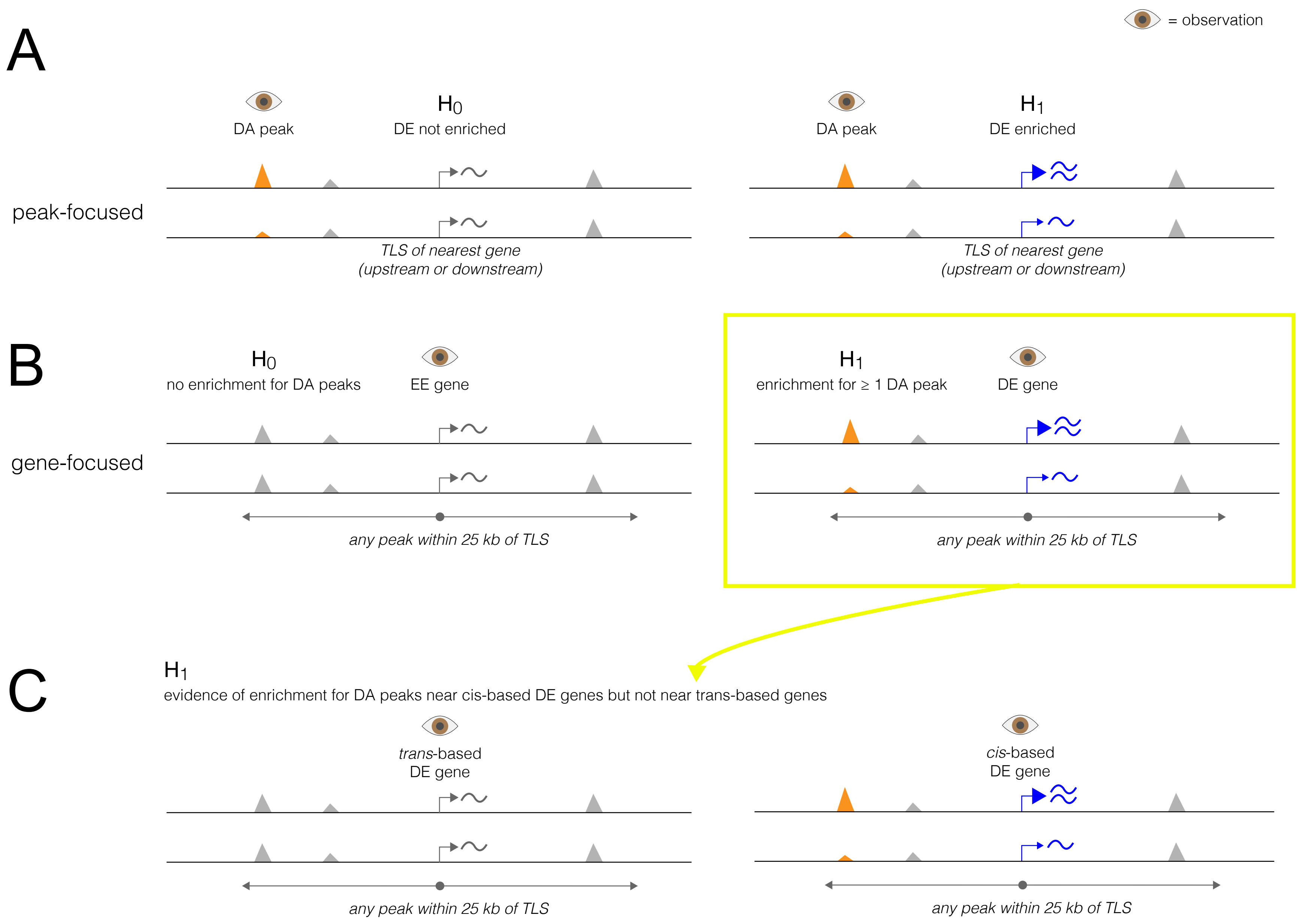

= observation
A
peak-focused
DA peak
H0
DE not enriched
TLS of nearest gene
(upstream or downstream)
DA peak
H1
DE enriched
TLS of nearest gene
(upstream or downstream)
B
gene-focused
H0
no enrichment for DA peaks
EE gene
any peak within 25 kb of TLS
H1
enrichment for ≥ 1 DA peak
DE gene
any peak within 25 kb of TLS
C
H1
evidence of enrichment for DA peaks near cis-based DE genes but not near trans-based genes
trans-based
DE gene
any peak within 25 kb of TLS
cis-based
DE gene
any peak within 25 kb of TLS

**Supplementary Figure 6.** Design of chi-squared tests for correlations among evolutionary changes in chromatin configuration and transcript abundance presented in Figure 5 (main text). In the following, DA = differentially accessible and DE = differentially expressed. **(A)** The "peaks-focused" test (Figure 5A). We considered all DA peaks, and asked if, for each peak, the nearest gene was DE. Under the null hypothesis $H_0$ (no connection between chromatin and transcription), we expect to see no enrichment for the nearest gene being DE. Under the alternative hypothesis $H_1$ (a connection between chromatin and transcription), we expect to see an enrichment for the nearest gene being DE. **(B)** The "genes-focused" test (Figure 5B). We considered all DE genes, and asked if at least 1 peak within 25 kb of that gene's translation start side was DA. Under the null hypothesis $H_0$ (no connection between chromatin and transcription), we expect to see no enrichment for DA peaks near that gene. Under the alternative hypothesis $H_1$ (a connection between chromatin and transcription), we expect to see an enrichment for DA peaks near that gene. **(C)** The "regulatory mode-focused" test (Figure 5C). For these tests, we began with the set of DE genes from Figure 5B) and subset them into those with expression differences based in *cis* or *trans*; we then asked whether *cis*- and/or *trans*-based DE genes had nearby DA peaks more often than expected by chance. The rationale is that *cis*-based DE is due to a local mutation that might act by altering chromatin configuration (although it could also act by altering transcription factor binding), whereas *trans*-based DE is not due to a local mutation and should not show any correlation with local chromatin. Under the null hypothesis $H_0$ (no connection between chromatin and transcription), we expect to see no enrichment for nearby DA peaks for either *cis*- or *trans*-based DE genes. Under the alternative hypothesis $H_1$ (a connection between chromatin and transcription), we expect to see an enrichment for nearby DA peaks for *cis*-based DE genes and possibly a depletion for nearby DA peaks for *trans*-based DE genes.
